# Reference-free clustering as an epidemiological tool for *Mycobacterium tuberculosis* lineage typing

**DOI:** 10.64898/2026.02.05.703994

**Authors:** Aureliana F. C. Chilengue, Daniel J. Whiley, Kate Cox, Maria Rosa Domingo-Sananes, Conor J. Meehan

## Abstract

Whole-genome sequencing (WGS) of *Mycobacterium tuberculosis* (Mtb) is widely used in the epidemiological investigation of recent transmission events, resulting in high-resolution strain typing. Accurate and rapid strain typing is essential for informing outbreak investigations and guiding tuberculosis control strategies. However, the gold-standard reference-guided SNP-calling pipeline currently used for strain typing relies on computationally intensive reference-mapping approaches, making it challenging to perform in many high-burden, resource-limited settings, where simplified and scalable genomic tools are urgently needed.

To address these limitations, we explored reference-free methods for medium resolution epidemiology, namely Mtb strain (lineage) typing, using a dataset of 535 complete genomes spanning the human- and animal-adapted lineages. Illumina paired-end reads were simulated from each complete genome, assembled, and analysed using three reference-free, k-mer-based tools: MASH, PopPUNK, and SKA2 (Split K-mer Analysis). Genetic distances were generated for each method and compared with a ground truth lineage assignment from with TB Profiler.

Our results demonstrated that reference-free methods can effectively distinguish Mtb lineages, with SKA2 showing the most promising performance across all datasets. SKA2 consistently recovered lineage and sub-lineage structure with high accuracy, demonstrating strong potential as an alternative to traditional WGS workflows. These findings highlight the utility of reference-free methods, particularly SKA2, for enabling accessible, scalable, and rapid Mtb strain typing, while supporting genomic epidemiology with low computational resources.

## 1. BACKGROUND

*Mycobacterium tuberculosis* (Mtb), the causative agent of tuberculosis (TB), is the leading cause of mortality globally due to pulmonary infection (WHO, 2023). According to the WHO (2023), Mtb infects nearly 100 million people annually, resulting in over 10.8 million active cases and 1.25 million deaths worldwide. The burden is disproportionately high in South-East Asia and Sub-Saharan Africa, where most cases are reported.

Mtb comprises 10 human-adapted lineages and several animal-adapted lineages, including La1-La3, *Mycobacterium microti*, and *Mycobacterium pinnipedii* (Goig *et al*., 2025). Each of these lineages exhibits unique epidemiological and phylogeographical characteristics (Goig *et al*., 2025), which are critical for understanding the global distribution and impact of TB. Lineages 2 and 4 are the most widespread, while lineage 1 causes the most infections in absolute numbers. Conversely, some lineages, such as 5-10, are more geographically restricted and often underrepresented in public datasets (Behruznia *et al*., 2025). Assigning lineage designations (often referred to as strain typing) to clinical strains is becoming increasingly important as evidence arises linking specific lineages to increased virulence and decreased drug susceptibilities (Orgeur and Brosch, 2018).

Recently, public health surveillance has been increasingly supported by molecular techniques to better understand the epidemiology of Mtb. The current gold standard for Mtb molecular epidemiology is whole-genome sequencing (WGS), which enables the identification of transmission clusters by detecting genetic variants such as single-nucleotide polymorphisms (SNPs) (Meehan *et al*., 2019). This approach also allows strain typing of human- and animal-adapted lineages, as well as their hierarchy of sub-lineages, through the analysis of variant calls and lineage-defining SNPs (Meehan *et al*., 2019). After SNPs are identified, strain typing is typically performed by comparing isolate-specific SNP patterns against a curated set of phylogenetically informative markers that reflect shared ancestry (Coll *et al*., 2014). These lineage-defining SNPs are used to classify isolates into hierarchical groups through phylogenetic reconstruction or barcode-based schemes, providing a medium-resolution and reproducible framework for characterising population structure and tracking evolutionary relationships (Coll *et al*., 2014; Homolka *et al*., 2012; Napier *et al*., 2020).

However, despite its strengths, WGS analysis relies on complex, computationally intensive pipelines to call SNPs, demanding significant bioinformatics skills and large computing resources (Spies *et al*., 2025). The gold standard for Mtb WGS is a mapping-based approach using the type strain H37Rv (a lineage 4 strain) as the reference. The dependence of the mapping-based approach on the H37Rv reference strain may introduce biases and limitations, as this reference does not adequately represent the global diversity of Mtb (Borrell *et al*., 2019). These requirements make WGS bioinformatics analysis challenging to perform in many endemic clinical settings (Rivière *et al*., 2021), especially when there is a need to deliver rapid results through simplified, low-computational-demand workflows that do not require highly trained bioinformaticians.

These limitations can be particularly challenging given the fact that the rapid identification of Mtb lineages is crucial for predicting potential future outbreaks (Napier *et al*., 2020) and for evaluating the effectiveness of disease control measures, including therapeutics and vaccines (Coll *et al*., 2014), whose efficacy may vary depending on the strain type (Bateson *et al*., 2022). Although mapping-based SNP-calling methods are widely regarded as the most valid approach for defining phylogenetic groupings with high confidence (Lipworth *et al*., 2019; Homolka *et al*., 2012; Coll *et al*., 2014), the comparably low level of genomic diversity in Mtb strains makes it challenging to achieve high-resolution discrimination within the complex (Homolka *et al*., 2012), particularly at finer phylogenetic scales (Coll *et al*., 2014).

These limitations highlight the need to complement mapping-based strategies with non-SNP-mapping methods, including reference-free approaches, to ensure scalable and consistent classification across diverse epidemiological contexts.

Reference-free methods have emerged as a promising alternative for epidemiology studies and have been widely used for strain typing and transmission analysis. These methods bypass the need for alignment or mapping to a set reference and undertake variant calling by directly comparing sequence content. As a result, they offer several advantages, including reduced computational complexity and processing time, and minimised reference bias, resulting in greater flexibility when analysing diverse strains (Lees *et al*., 2019; Zielezinski *et al*., 2017).

Such approaches have the potential to capture a more complete picture of genetic diversity, including in genomic regions that might be missed by reference-based approaches (Zielezinski *et al*., 2017). Examples include k-mer based methods such as MASH (Ondov *et al*., 2016) and PopPUNK (Lees *et al*., 2019), which estimate genetic distances between isolates using shared k-mer content. Another tool, SKA2 (Derelle *et al*., 2024), uses split k-mers to identify SNPs between genomes without the need for alignment. These tools have shown variable accuracy for Mtb in the past ranging from low correlation for transmission detection (Lees *et al*., 2019) to similar results compared to the gold standard (Derelle *et al*., 2024), albeit all on simulated data. This study aims to explore the accuracy of these reference-free tools for rapid and accurate strain typing of Mtb, specifically the identification of Mtb (sub-)lineages as a way to enable more efficient and accessible genomic epidemiology in several settings (Zielezinski *et al*., 2019), particularly in low-resource environments.

## 2. METHODS

### 2.1 Dataset

#### Selection and characterisation of the genomes

A total of 535 complete and closed (single contig) genome assemblies from human-adapted (L1-L9) and animal-adapted (La1, La2, La3, *M. microti* and *M. pinnipedii*) lineages across Mtb were selected for this study (Supplementary Table 1). This dataset consisted of those from a previously published study (Behruznia *et al*., 2025), which collected all closed genomes from NCBI up until 2023. The same procedure was repeated to complement this collection with all complete Mtb genomes from NCBI up until April 2025. Quality assessment of the assemblies was performed using BUSCO v5.4.7 (Manni *et al*., 2021) to ensure only those with ≥ 95% completeness were included in the study. Lineage assignment was done using TB-profiler version 5.2 (Phelan *et al*., 2019) and was defined as the ground truth assignments. Spoligotype patterns were also estimated for each isolate from its genome using the --spoligotype option in TB-profiler version 5.2 (Napier *et al*., 2020).

#### Simulated Reads

Paired-end Illumina-like reads (300 bp) were simulated from 535 complete Mtb genomes using InSilicoSeq v1.5.4 (Gourlé *et al*., 2019) to a depth of ∼100X. The simulation employed the built-in MiSeq error model, which reproduces empirical sequencing features such as substitution, insertion and deletion errors, GC bias, and quality score distributions. This model is derived from aligned reads in the PRJEB20178 dataset and accurately mimics Illumina MiSeq sequencing characteristics (Gourlé *et al*., 2019). The simulated reads were output as compressed FASTQ files for each genome and used for further analysis.

#### Assembly of Simulated Reads

The simulated paired-end reads were assembled using SKESA v2.4.0 (Souvorov, Agarwala and Lipman, 2018), a conservative *de novo* assembler optimised for Illumina sequencing data. For each genome, paired FASTQ files (*R1.fastq.gz and *R2.fastq.gz) were provided to SKESA using default parameters. The resulting assemblies were saved in FASTA format and used for downstream analyses.

### 2.2 SNP distance estimations using DNAdiff

The 535 genome assemblies were compared pairwise using DNAdiff v1.3 from the MUMmer4 tool suite (Marçais *et al*., 2018). The resulting GSNP counts, where each SNP is bound by 20 exact matches to reduce false positives, were then used to compute pairwise SNP distances between isolates.

### 2.3 Reference-free methods for clustering genomes

Three reference-free methods were employed in our analysis: PopPUNK (Lees *et al*., 2019), MASH (Ondov *et al*., 2016), and SKA2 (Derelle *et al*., 2024). These methods were applied to complete genomes, simulated reads, and assembled genomes to assess their consistency across different data types.

We ran PopPUNK version 2.6.0 (Lees *et al*., 2019) using the ***poppunk --create-db*** command with a minimum k-mer size of 29, maximum k-mer size of 61, and a sketch size of 100,000 to generate compact sketches of each genome. These sketches are then compared using the MinHash algorithm to efficiently estimate core and accessory genome distances based on shared k-mer content, which was used to cluster the genomes and define population structures according to genome similarity across the dataset. The core and accessory distances were subsequently extracted using the ***poppunk_extract_distances*.*py*** command.

Since MASH (Ondov *et al*., 2016) operates using a single k-mer size, we ran the ***mash sketch*** (version 2.3) command with a k-mer size of 31 and a sketch size of 100,000 to generate the genome sketches. These sketches are then compared using the MinHash algorithm as above. The resulting pairwise distances were extracted using the ***mash dist*** command and used to cluster the genomes according to their genetic similarities.

For SKA2 (Derelle *et al*., 2024) version 0.4.0, we used the ***ska build*** command with a k-mer size of 31 to generate the split k-mers with a variable middle base. These split k-mers were used as keys in a hash map, with their middle bases stored as values. After building the dictionaries for all genomes, the genetic distances between all pairs of genomes were calculated using the ***ska distance*** command by iterating over every pair of samples to compare their split k-mer dictionaries. For each shared split k-mer, SKA2 assessed whether the middle nucleotide matched between genomes by comparing the nucleotide stated at that position..

### 2.4 Genetic Distance Distribution

To compare the genetic distance distributions within and between lineage groupings, we used the distance matrices generated by each method (DNAdiff, PopPUNK, MASH, and SKA2) and combined them with the TB-Profiler-defined lineage assignments. Using a custom R script, we matched each genome pair to its corresponding lineage classification, labelling pairs as “within-lineage” or “between-lineage,” and assigned them to the appropriate lineage group or the “between” category. We then visualised these distance distributions using histograms and density plots with the ggplot2 package (Wickham, 2016), applying custom colour schemes to highlight lineage-specific patterns. This approach enabled us to evaluate how each method addressed genetic distances and whether it effectively maintained separation between distinct phylogenetic groups.

All scripts for this analysis can be found in the GitHub repository https://github.com/Aurelianachilengue/mtb-strain-typing.

### 2.5 Reference-Free Clustering and Lineage Delineation

Using the genetic distance matrices generated from the four methods, we built trees using hierarchical clustering in R (version 4.3.2) using the APE package (version 5.7.1) (Paradis and Schliep, 2019). These trees were then visualised using ITOL-Interactive Tree of Life (Letunic and Bork, 2021) to illustrate the separation and delineation of lineages for each method.

### 2.6 Cluster membership accuracy

A lineage cluster membership accuracy ratio was used to evaluate how accurately each method categorised the isolates into clusters corresponding to the TB-profiler lineage designations. Since lineages are monophyletic groups, we used a phylogenetic clustering approach to determine if lineages formed clades in the dendrograms constructed by each of the SNP distance estimation approaches. This was conducted to assess the ability of each approach to accurately resolve a given lineage-based cluster.

For this analysis, we used a custom Python script that took a tree file in Newick format and a file containing all the isolate names of a given lineage. This methodology is illustrated in Supplementary Figure S1. For a given lineage-based cluster (as determined by TB-profiler), the most recent common ancestor (MRCA) node was determined within the tree and the subtree that contained all the taxa that descended from that MRCA was extracted from the parent tree. The number of taxa in the original lineage cluster and the number of taxa in the extracted subtree were calculated. Subsequently, the total count of isolates in each lineage was divided by the subtree count from each method. If these two sets were identical (i.e. the subtree of the cluster contained only taxa that were designated to be in that lineage-based cluster), the method accurately grouped that lineage, giving a ratio of 1. A ratio below 1 reflected either misclassification, where isolates assigned to one sub-lineage were grouped into a different sub-lineage, or extended clustering, where isolates without sub-lineage designation were grouped into a specific sub-lineage cluster.

To identify the isolates being grouped incorrectly by each method (i.e. a clustering ratio <1), we coloured the subtree based on lineage assignment (Supplementary Figure S1). This allowed for visual inspection of isolate clustering and identification of the cause of over-clustering. Spoligotype patterns were then used to further explain any unexpected grouping of these genomes.

Genomes lacking sub-lineage designations were only labelled at the higher lineage level. Therefore, we evaluated clustering based on the available designations and did not assess separation at finer levels where assignments were missing.

## RESULTS

Our dataset comprised 535 closed Mtb genomes covering human-adapted and animal-associated lineages. Lineage assignment was performed using TB-Profiler, which served as the ground truth for reference-free analyses. Pairwise whole-genome SNP distances were computed using DNAdiff for genome-to-genome comparisons. Alongside this, three k-mer-based, reference-free methods (PopPUNK, MASH, and SKA2) were systematically analysed in concordance with TB-Profiler lineage assignments using as input the complete genomes, simulated reads or re-assembled genomes.

### Genetic distance distributions across Mtb lineages using reference-free approaches

We used the genetic distance distributions within and between lineages based on the various methods and datasets to assess their robustness at differentiating lineages from each other. For the complete genomes, DNAdiff and SKA2 SNP-based distances produced the strongest separation between intra-lineage and inter-lineage distances (Supplementary Figure S2, Figure1). In contrast, PopPUNK and MASH k-mer-based distances showed broader intra-lineage distributions and greater overlap with inter-lineage distributions of genetic distances, with MASH exhibiting the most pronounced overlap.

When applied to the simulated reads, SKA2 produced well-separated intra- and inter-lineage distance distributions, while PopPUNK and MASH showed overlap between the two (Supplementary Figure S3, Figure 1). After assembling the simulated reads, PopPUNK and MASH regained the expected separation, and SKA2 showed consistent distance patterns across all data types (Supplementary Figure S4, Figure 1).

**Figure 1.**
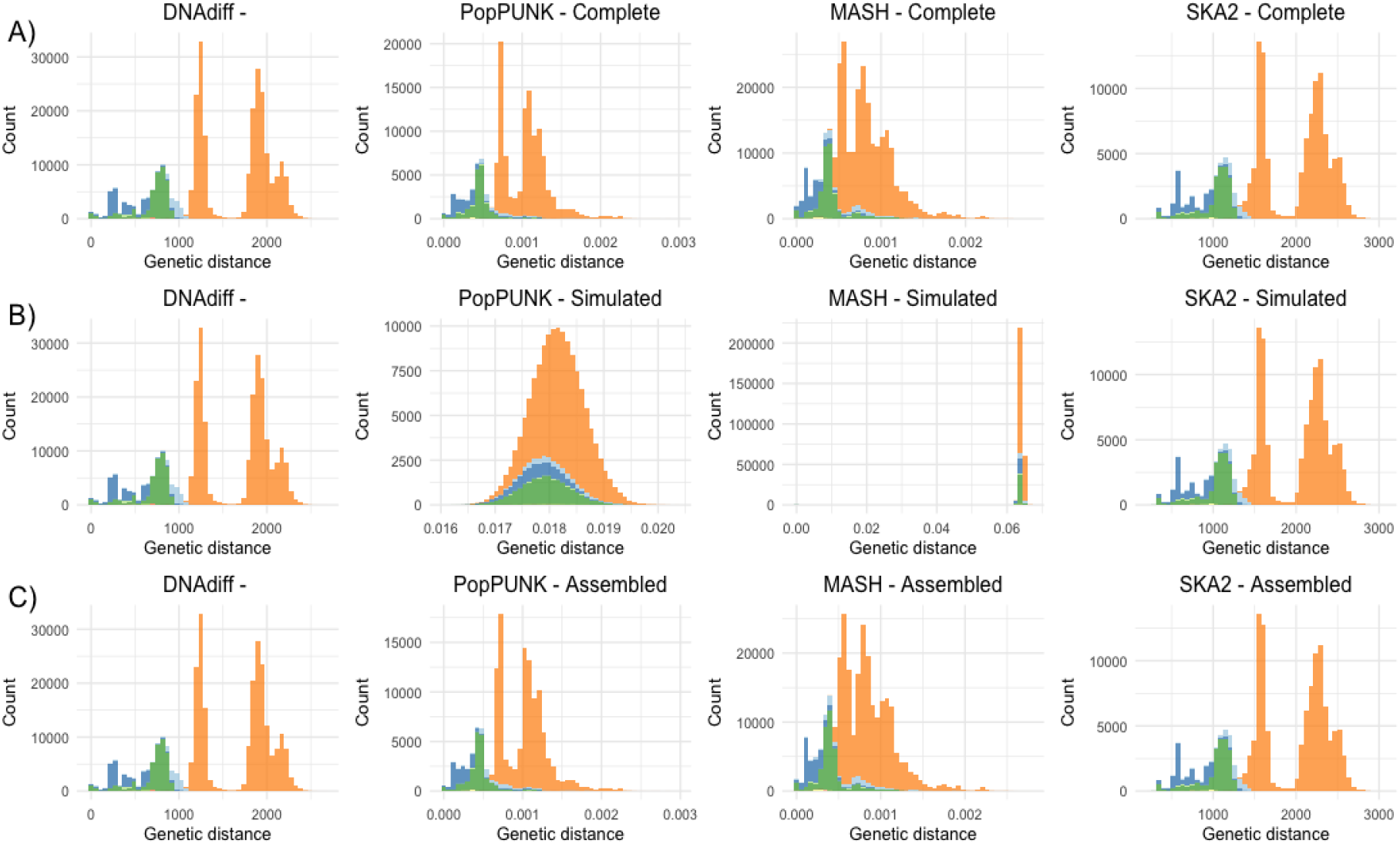
Comparative distribution of genetic distances within and between Mtb lineages using complete genomes (A), simulated reads (B), and assembled simulated reads (C). Inter-lineage distances are shown in orange, while intra-lineage distances are coloured by lineage for each method. DNAdiff used complete genomes in all analyses.

### Reference-free tools for delineating the lineages of Mtb

All three reference-free methods (PopPUNK, MASH, and SKA2) recovered the same major grouping structure as the TB-Profiler lineage classifications when given complete genomes as input (Lineages 1-9 and animal-associated lineages La1, La2, La3, M. microti, and *M. pinnipedii*). The resulting hierarchical clustering trees matched the expected grouping patterns based on the full-genome DNAdiff distance-based clustering (Supplementary Figure S5). A visual comparison of the clustering dendrograms revealed broadly similar topologies across the methods. Minor differences in branch resolution and cluster compactness were observed for PopPUNK and MASH, whereas SKA2 showed clear separation between lineage groupings. All methods produced stable and coherent grouping behaviour when run on the complete genomes.

Using the raw simulated reads as input, SKA2 maintained clear lineage-level grouping separation, while PopPUNK and MASH did not consistently recover several lineage groupings (Supplementary Figure S6). After assembling the simulated reads, PopPUNK and MASH produced lineage groupings similar to those observed with the complete genomes, and SKA2 consistently showed strong lineage-level grouping across all data types (Supplementary Figure S7).

### Reference-free tools reveal varying performance in clustering Mtb genomes across lineage levels

For the complete genomes included in this study, all three reference-free methods achieved perfect classification at the lineage level, with a cluster membership ratio of 1.0 (Supplementary Figure S8) across all major Mtb lineages, demonstrating alignment with DNAdiff and correctly grouping genomes according to TB-profiler lineage assignment.

At the sub-lineage level, some notable deviations were observed in groupings. Approximately 62% of the genomes broadly assigned to Lineage 3 and lacking sub-lineage designation were grouped into sub-lineage L3.1 by all three reference-free methods and DNAdiff, with a consistent drop in the ratio below 1.0 (Supplementary Figure S9, Figure 2). Spoligotype analysis showed that both the L3 and L3.1 genomes in this cluster shared the CAS1 pattern.

**Figure 2.**
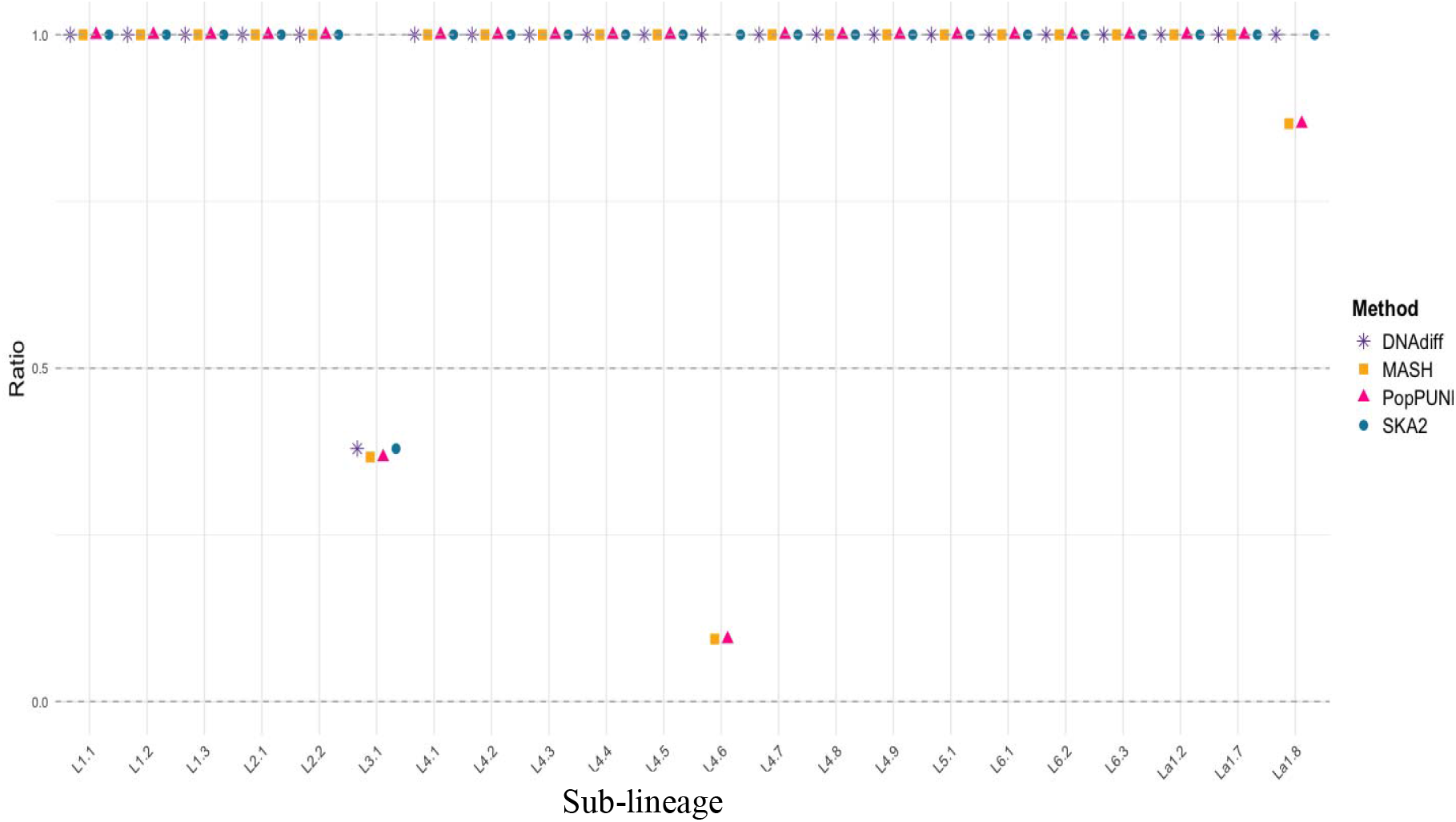
Lineage cluster membership accuracy ratio for each method using complete genomes. A ratio of 1.0 indicates perfect agreement between each method’s clustering results and the TB-Profiler–assigned lineage groups. Ratios below 1.0 reflect over-clustering, where genomes from different lineage groups were incorrectly merged into the same cluster. A total of 494 genomes were included in this analysis.

Similarly, 90.6% of isolates from sub-lineages L4.8, L4.9, and L4.7 were grouped within L4.6 by PopPUNK and MASH, and all these genomes displayed LAM spoligotype patterns. Additionally, 13.3% of La1.7 isolates clustered with La1.8, consistent with the shared BOV-family spoligotype pattern (Supplementary Figure S9, Figure 2).

The clustering performance at finer lineage resolutions (e.g., sub-sub-lineage) was generally accurate for SKA2 and consistent with DNAdiff and TB-profiler. However, 85.7% of genomes originally assigned to Lineage 3 were grouped within the sub-sub-lineage L3.1.2, across all methods, including SKA2 and DNAdiff (Supplementary Figure S10). A similar pattern was seen for L3.1.1, where 85.7% of Lineage 3 genomes were clustered into this group by PopPUNK and MASH. These two methods also grouped genomes originally assigned to L1.1 into L1.1.3, while 37.5% of genomes from La1.8.1 were clustered with La1.7. At the sub-sub-sub-lineage level, all methods maintained perfect clustering (ratio = 1.0) across all groups (Supplementary Figure S11), although only 29% (157/535) genomes had a designation at this level.

When applied to simulated reads, PopPUNK and MASH showed consistently poor clustering performance across all lineage levels, with cluster membership ratios falling below 1.0 for most lineages, particularly for PopPUNK (Supplementary Figure S12-S15, Figure 3). At all lineage levels, genomes from different lineages were frequently grouped together, with inter-lineage mixing. At the lineage level, correct clustering was observed only for L7, which included two genomes, for both PopPUNK and MASH. At finer lineage resolutions, correct clustering by MASH was observed only in a small number of cases involving very limited group sizes, including sub-sub-sub-lineages L1.2.1.1, L4.1.1.1, and L4.2.2.2 (each comprising two genomes) (Supplementary Figure S15; Figure 3), whereas PopPUNK did not cluster correctly at these finer resolutions. Following genome assembly, clustering performance for both PopPUNK and Mash improved, with results matching those obtained from complete genome sequences (Supplementary Figure S16-S19). In contrast, SKA2 produced identical results for simulated reads, assembled genomes, and complete genomes.

**Figure 3.**
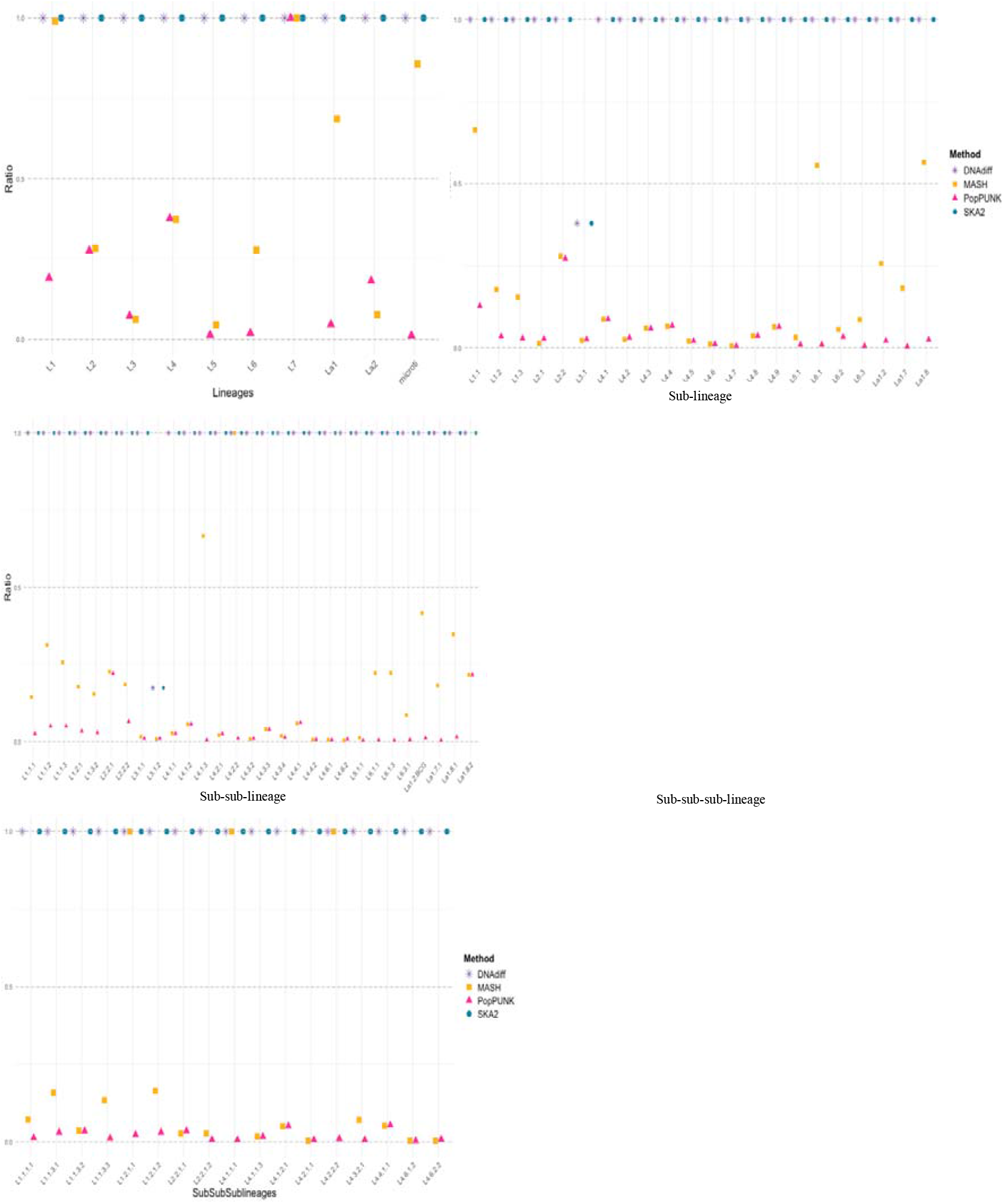
Lineage cluster membership accuracy ratio for each method using simulated reads. A ratio of 1.0 indicates perfect agreement between each method’s clustering results and the TB-Profiler–assigned lineage groups. Ratios below 1.0 reflect over-clustering, where genomes from different lineage groups were incorrectly merged into the same cluster. A total of 494 genomes were included in this analysis.

## DISCUSSION

Genetic distance comparisons enable the evaluation of how effectively each method separates isolates within and between Mtb lineages. For the complete genomes, SKA2 produced the clearest separation between intra- and inter-lineage distances, closest to DNAdiff (Figure 1, Supplementary Figure S2) and more concordant with TB-profiler. In contrast, PopPUNK and MASH showed broader within-lineage distributions with noticeable overlap in the inter-lineage range, particularly for MASH. In the simulated read dataset, SKA2 maintained strong separation, while PopPUNK and MASH displayed extensive overlap between intra- and inter-lineage distances (Figure 1, Supplementary Figure S3), indicating reduced discriminatory power when applied directly to unassembled reads. After assembling the simulated reads, PopPUNK and MASH regained the expected lineage-level separation (Figure 1, Supplementary Figure S4), more closely resembling the patterns observed for complete genomes, and SKA2 remained highly accurate and consistent across the different input dataset.

These results suggest that SKA2’s split k-mer approach is more robust to diverse input data types, whereas PopPUNK and MASH are more sensitive to simulated read-level noise…

Nevertheless, the residual overlap observed across all methods underscores the continued challenge of achieving high-resolution clustering using reference-free approaches, particularly in a genetically homogeneous pathogen like *M. tuberculosis*. This has important implications if such methods are to be implemented for recent transmission analyses, where the ability to resolve closely related isolates is critical for accurately inferring transmission events (Martin *et al*., 2018; Walker *et al*., 2013).

The hierarchical clustering trees generated by PopPUNK, MASH, and SKA2 demonstrated that reference-free methods can effectively capture and delineate the broad population structure of *M. tuberculosis*, including human and animal-adapted lineages. All three methods successfully reproduced the lineage groupings defined by TB-Profiler and those obtained through DNAdiff-based genome comparisons (Supplementary Figure S5). These findings support previous studies showing that reference-free k-mer-based approaches can approximate phylogenetic relationships in bacterial populations (Lees *et al*., 2019; Gardner and Hall, 2013). Despite their shared ability to resolve the major Mtb lineages, the resulting tree topologies revealed differences in branch lengths and resolution. Based on our results, SKA2 produced branching patterns more consistent with DNAdiff distances calculated from whole-genome alignments, recovering the major lineages as distinct clades and suggesting a higher resolution in capturing evolutionary relationships (Supplementary Figure S5).

The low resolution observed for PopPUNK and MASH in delineating several lineage groupings from simulated reads (Supplementary Figure S6) likely reflects the known limitations of applying k-mer-based methods with heuristic distance calculations directly to raw sequencing data. Raw reads can be more fragmented and often contain sequencing errors, which likely introduce false k-mers and reduce the accuracy of distance estimation (Lees *et al*., 2019; Ondov *et al*., 2016). Although both PopPUNK and MASH can run on unassembled reads, previous studies have reported that the best-hit strain is often incorrect due to noise in the raw data, and that assembly or error-filtering significantly improves resolution (Lees *et al*., 2019; Ondov *et al*., 2016). After assembly with SKESA, both PopPUNK and MASH recovered the expected lineage grouping structure (Supplementary Figure S7), consistent with previous findings that assembly enhances the reliability of k-mer-based clustering analyses (Lees *et al*., 2019; Ondov *et al*., 2016; Souvorov, Agarwala and Lipman, 2018).

We observed variation in how DNAdiff and the reference-free methods classified genomes into clusters corresponding to the lineage groups assigned by TB-Profiler when using complete genomes. Nevertheless, all methods showed consistent agreement with TB-Profiler-defined lineages at the broad lineage level (Supplementary Figure S8), highlighting both the reliability of reference-free approaches for broad lineage classification and the robustness of the SNP markers employed by TB-Profiler. The strong performance of these k-mer-based methods at this resolution highlights their practical relevance for public health surveillance, where rapid and accurate lineage identification is essential for informing outbreak investigations and guiding TB control strategies (Coll *et al*., 2014; Couvin *et al*., 2025).

All methods showed a pattern of extended clustering when grouping isolates from sub-lineage L3.1 (Supplementary Figure S9); a group historically recognised for its poorly defined phylogenetic boundaries (Shuaib *et al*., 2020; Xu *et al*., 2022). Notably, when using TB-Profiler as the classification baseline, several genomes assigned to L3.1 by DNAdiff and the reference-free tools were originally classified only as Lineage 3, without sub-lineage designation. This pattern likely reflects the intrinsic complexity of defining sub-lineage L3.1, as Lineage 3 is known to lack clear resolution at this level. For instance, a study from eastern Sudan also reported similar challenges, noting that although L3 was the predominant lineage, most isolates could not be confidently assigned to specific sub-lineages, highlighting current limitations in the sub-lineage classification of L3 (Shuaib *et al*., 2020). Likewise, a study from the Kashgar Prefecture (China) reported the identification of four previously unclassified clades within Lineage 3, suggesting that the current sub-lineage framework does not fully encompass the genetic diversity of L3 strains in the region (Xu *et al*., 2022).

These findings collectively highlight the need to refine current sub-lineage classifications to better reflect region-specific diversity within Lineage 3. Furthermore, the observed extended clustering may suggest that these methods are capturing additional population structure not currently resolved by SNP-based classification schemes, potentially reclassifying some Lineage 3 strains as L3.1. Future investigations could explore whether specific SNPs show complete differentiation between the groups defined by the reference-free methods (i.e., fixation index, F_ST_ = 1), providing a basis for improving and refining existing sub-lineage designations.

Beyond the challenges observed in L3.1, misclassification was identified within sub-lineage 4, where PopPUNK and MASH grouped genomes from sub-lineages L4.8, L4.9, and L4.7 into L4.6 (Supplementary Figure S9). This pattern suggests that these sub-lineages share a high degree of genetic similarity, potentially exceeding the resolving power of current sub-lineage definitions. The reduced discriminatory power when differentiating groups with minimal genomic divergence further highlights the broader challenge of achieving fine-scale resolution in *M. tuberculosis*, as previously described by (Coll *et al*., 2014). Consistent with this observation, all these genomes displayed LAM spoligotype patterns, indicating a shared ancestral background that may obscure finer phylogenetic distinctions. Previous genomic studies described weak separation among several L4 sub-lineages and noted that many globally distributed Lineage 4 groups exhibit overlapping population structures (Stucki *et al*., 2016). Additionally, Sabin et al. (2020) reported that sub-lineages L4.7, L4.8, and L4.9 fall within the broader L4.10/PGG3 clade, a grouping originally defined by Stucki et al. (2016), suggesting that their current subdivision may not accurately reflect true phylogenetic differentiation. Taken together, these findings imply that the grouping of these genomes by k-mer-based methods likely reflects genuine genomic similarity rather than methodological artefacts. Such results highlight that some apparent inconsistencies may arise from limitations in the existing taxonomic framework, emphasising the need to re-evaluate sub-lineage boundaries within Lineage 4 to ensure accurate representation of its evolutionary diversity. Notably, SKA2 maintained strong discriminatory performance across all other evaluated sub-lineages, reaffirming its robustness as a reference-free tool for fine-scale lineage classification.

A similar pattern of apparent misclassification was initially observed for some isolates previously designated as L1.1 and L1.2, which were grouped together by all methods, including DNAdiff. However, this discrepancy was later resolved following the updated redefinition of Mtb Lineage 1 sub-lineages, where the former L1.2.2 was reclassified as L1.3 (Netikul *et al*., 2022). This revision aligned the clustering patterns observed across PopPUNK, MASH, SKA2, and DNAdiff with the revised lineage framework, confirming that the apparent inconsistency stemmed from outdated lineage definitions rather than methodological inaccuracies. This observation underscores the dynamic nature of Mtb taxonomy and highlights the importance of continuous refinement of reference databases and lineage nomenclature to ensure accurate genomic classification.

Interestingly, at the sub-sub-sub-lineage level, all methods achieved perfect clustering concordance (Supplementary Figure S11), suggesting strong discriminatory performance at this fine scale. However, it is important to note that only 151 genomes were assigned at this resolution, and most major Mtb lineages were not represented at this level. This limited diversity and reduced number of assignments may have simplified the clustering task, thereby inflating the apparent concordance. This may represent the practical limit of lineage subdivision, beyond which further resolution may reflect transmission clusters.

Beyond clustering accuracy, practical considerations such as computational efficiency and ease of deployment are central to determining whether reference-free methods can be adopted in routine surveillance and clinical workflows. In this study, SKA2 completed analyses within approximately 20 minutes using ∼300 MB of memory, demonstrating that it can be executed on standard laboratory computers without access to high-performance computing infrastructure. Together, these characteristics highlight the utility of reference-free methods, particularly SKA2, for enabling accessible, scalable, and rapid Mtb strain typing while supporting genomic epidemiology in low-resource environments. Overall, our findings reinforce the potential of reference-free approaches for strain typing and lineage assignment in Mtb, particularly when reference-based tools are not feasible, such as during rapid outbreak investigations or in resource-limited settings where timely identification of circulating strains is essential. However, differences in resolution capacity between tools should be carefully considered when selecting a method for downstream epidemiological or evolutionary analyses.

In this study, we systematically evaluated the performance of three reference-free clustering methods PopPUNK, MASH, and SKA2, against the TB-profiler defined lineages and DNAdiff-derived lineage groupings to assess their applicability for strain typing and cluster definition. While all methods showed strong concordance at the lineage level, finer-scale resolution varied, reflecting both methodological and biological constraints. While these findings highlight the promise of reference-free approaches for rapid lineage-level classification and surveillance, some limitations should be acknowledged. The reliance on current SNP-based definitions, where some genomes lack sub-lineage designations, may limit resolution and overlook subtle or emerging diversity, underscoring the need for continued refinement of genomic markers and classification frameworks. Future work should expand evaluations to include geographically diverse collections, and routine sequence reads, beyond simulated data, to explore how reference-free approaches can support not only lineage delineation but also recent transmission analyses in real-world epidemiological contexts.

## Author Notes

All supporting data, codes have been provided within the article or through supplementary data files. Nineteen supplementary figures and one supplementary table are available in the online Supplementary Material.

## Abbreviations

Mtb: *Mycobacterium tuberculosis*
TB: tuberculosis
WGS: whole-genome sequencing
SNP: single-nucleotide polymorphism
PopPUNK: Population Partitioning Using Nucleotide K-mers
MASH: MinHash-based sequence distance estimation; SKA2
SKA2: Split K-mer Analysis
DNAdiff: genome comparison tool from the MUMmer package

## Impact Statement

Tuberculosis (TB) remains one of the leading causes of mortality from infectious diseases worldwide. Over the past decade, whole-genome sequencing (WGS) has been increasingly used to support surveillance and outbreak investigations by enabling high-resolution strain typing to inform disease control efforts. However, the gold-standard reference-guided SNP-calling pipelines used for WGS-based strain typing are computationally intensive, limiting their routine use in many resource-limited, high-burden settings where rapid and scalable strain typing is essential for effective TB control. In this study, we evaluate reference-free tools for *Mycobacterium tuberculosis* (Mtb) strain typing using a large and diverse collection of complete genomes and simulated sequencing reads for each complete genome. We show that reference-free tools can reliably recover lineage structure, with approaches such as SKA2 performing efficiently across datasets while remaining computationally efficient and easy to deploy. This work highlights the potential of reference-free to enable accessible, scalable, and rapid strain typing, supporting broader adoption of genomic epidemiology in low-resource settings.

## Data Summary

All complete Mtb genomes analysed in this study were retrieved from the NCBI RefSeq and GenBank databases, including genomes previously reported by Behruznia *et al*. (2025) and additional complete genomes available up to April 2025. Simulated Illumina paired-end reads were generated from these genomes using InSilicoSeq v1.5.4. Accession numbers for all genomes included in the analysis are provided in Supplementary Table 1.

## Author contributions

A.F.C.C contributed to conceptualization, methodology, formal analysis, investigation, data curation, writing -original draft, writing – reviewing & editing, visualization and funding acquisition.

D.J.W. contributed to conceptualization, methodology, formal analysis, data curation, writing– reviewing & editing.

K.C. contributed to methodology, formal analysis, writing – reviewing & editing.

M.R.D.S. contributed to conceptualization, methodology, formal analysis, investigation, writing -original draft, writing – reviewing & editing, supervision, and visualization.

C.J.M. contributed to conceptualization, methodology, formal analysis, investigation, writing-original draft, writing – reviewing & editing, supervision, project administration and funding acquisition.

