## Supplementary Figure for "Reference-free clustering as an epidemiological tool for *Mycobacterium tuberculosis* lineage typing"

### Supplementary Figures

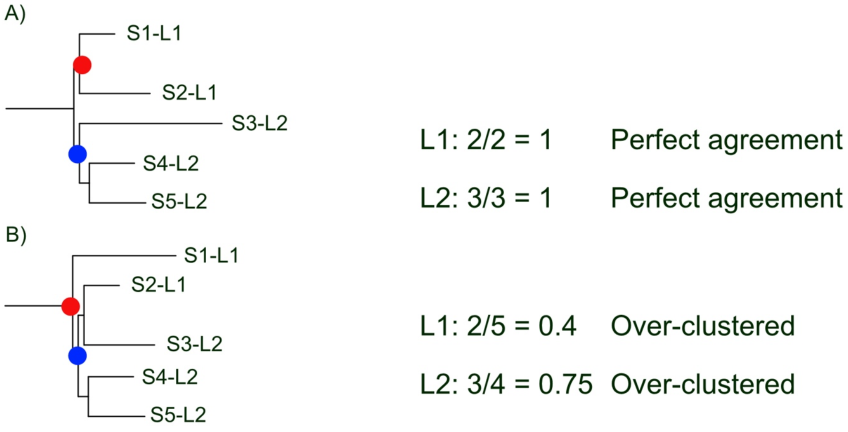


**Figure S1**. Illustration of lineage assignment agreement calculation. A lineage is assigned to each sample by TB-Profiler (indicated by the -L1 or -L2 in the sample name). A reference-free approach calculates distances between the samples and a dendrogram is created from that distance. The MRCA of each lineage (based on TB-profiler) is determined in the dendrogram (indicated by the red dot for L1 and blue dot for L2). The number of samples in that lineage is divided by the number descended from that MRCA to look for agreement. In A), the dendrogram separates that lineages properly, so perfect agreement is calculated, giving a score of 1. In B), the dendrogram incorrectly places one L1 sample in a clade with L2 samples, increasing the number of samples descended from each MRCA of each lineage, resulting in incorrect clustering.

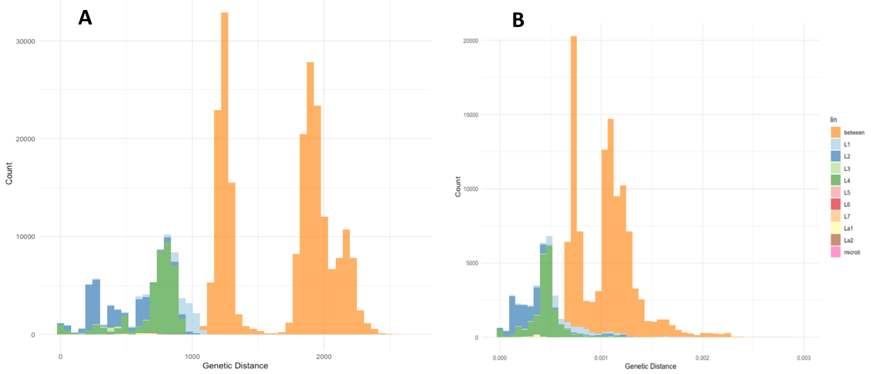

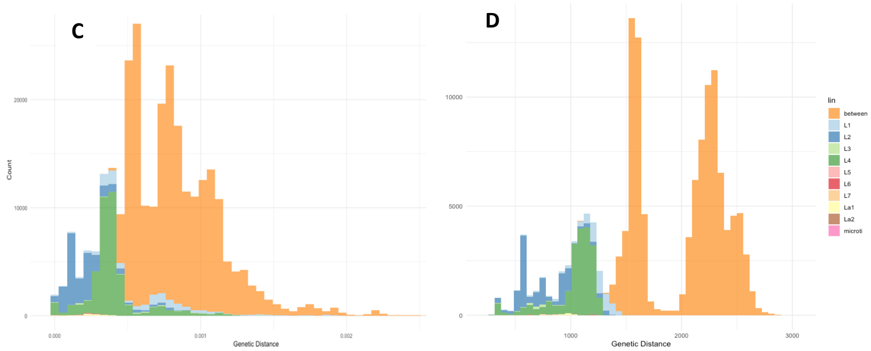


**Figure S2**. Comparative distribution of genetic distances within and between *M. tuberculosis* lineages (complete genomes). Distances between lineages are shown in orange, while within-lineage distances are coloured by lineage for each method: **DNAdiff (A), PopPUNK (B), MASH (C), SKA2 (D)**.

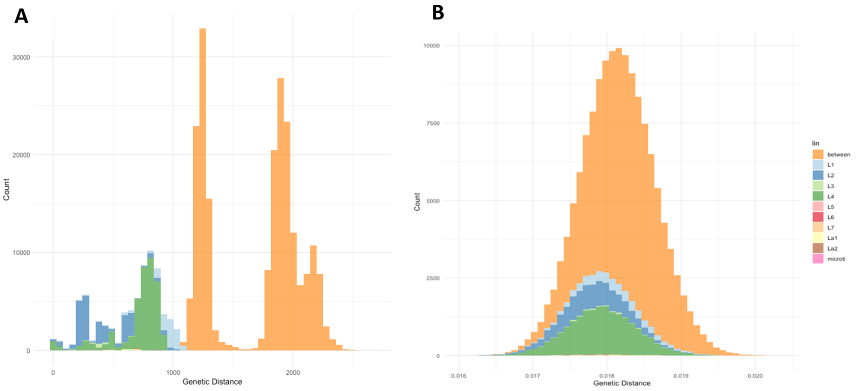

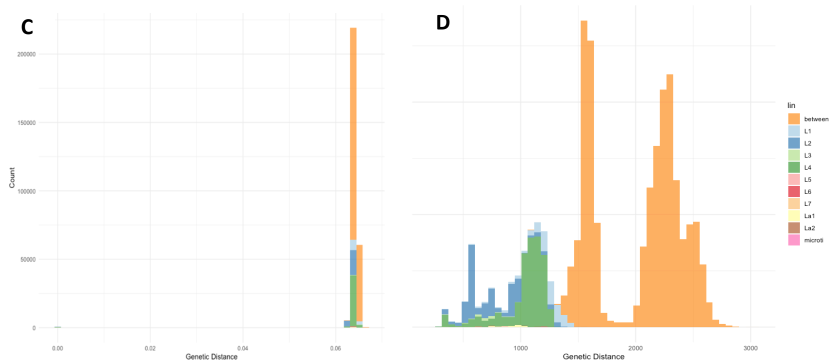


**Figure S3.** Comparative distribution of genetic distances within and between *M. tuberculosis* lineages (simulated reads). Distances between lineages are shown in orange, while within-lineage distances are coloured by lineage for each method: **DNAdiff using complete genomes (A), PopPUNK (B), MASH (C), SKA2 (D)**.

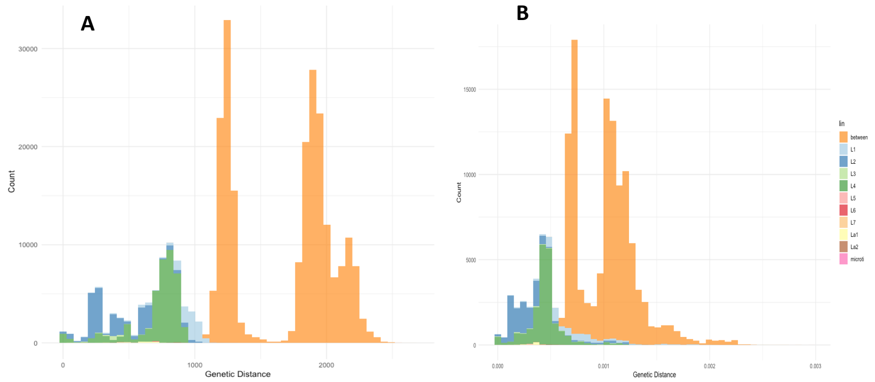

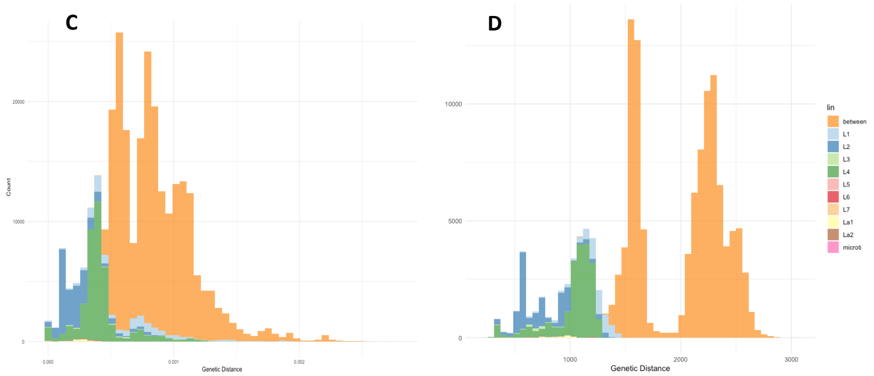


**Figure S4.** Comparative distribution of genetic distances within and between *M. tuberculosis* lineages (assembled simulated reads). Distances between lineages are shown in orange, while within-lineage distances are coloured by lineage for each method: **DNAdiff (A), PopPUNK (B), MASH (C), SKA2 (D)**.

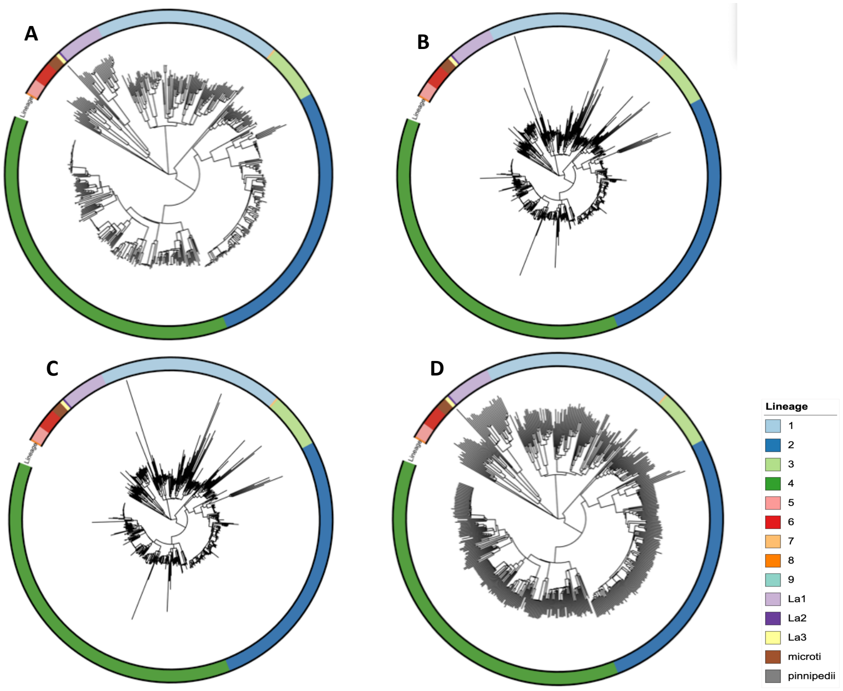


**Figure S5**. Delineation of lineages and relationships among the complete genomes. Hierarchical clustering trees constructed from distance matrices generated by **DNAdiff (A)**, **PopPUNK (B)**, **MASH (C)**, and **SKA2 (D)**. All trees are rooted on the Lineage 8 strain**.**

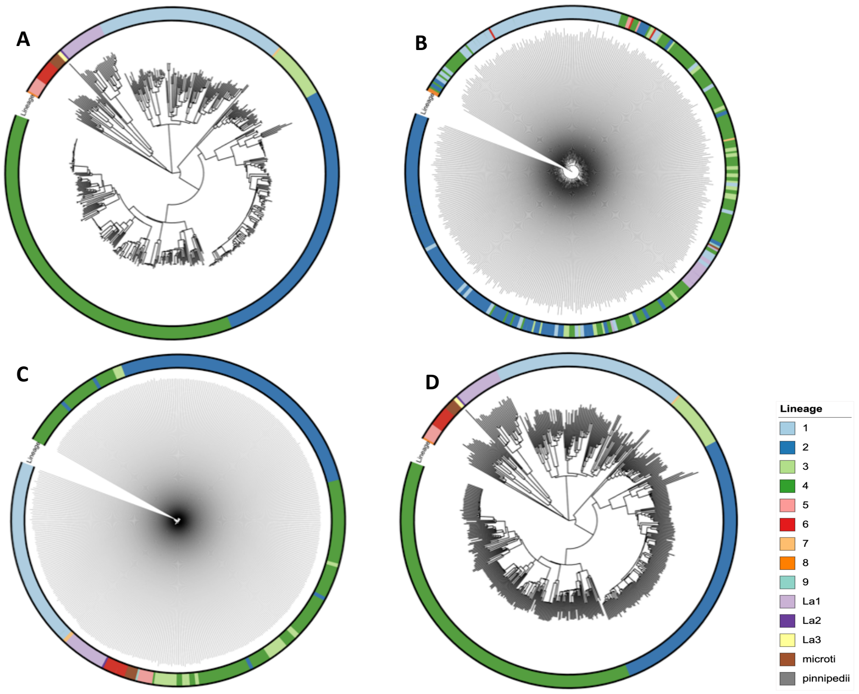


**Figure S6**. Delineation of lineages and relationships among the simulated reads. Hierarchical clustering trees constructed from distance matrices generated by **DNAdiff (A)**, **PopPUNK (B)**, **MASH (C)**, and **SKA2 (D)**. All trees are rooted on the Lineage 8 strain.

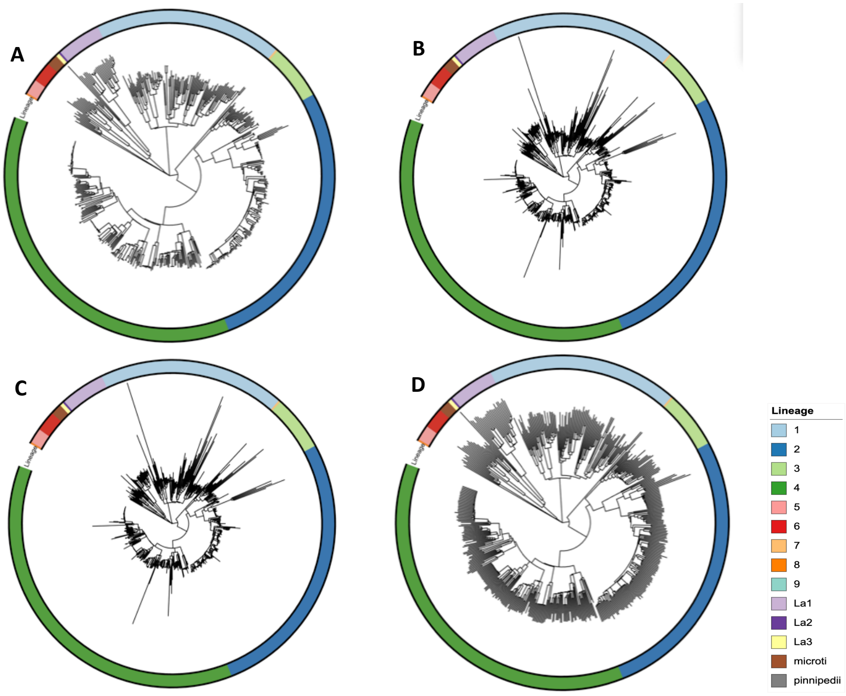


**Figure S7**: Delineation of lineages and relationships among the assembled simulated reads. Hierarchical clustering trees constructed from distance matrices generated by **DNAdiff (A)**, **PopPUNK (B)**, **MASH (C)**, and **SKA2 (D)**. All trees are rooted on the **Lineage 8 strain**.

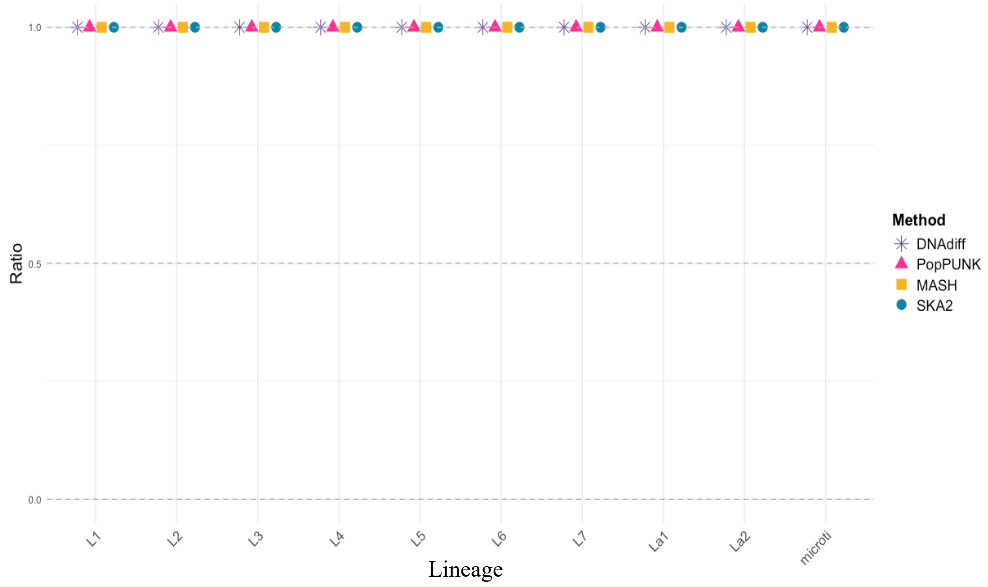


**Figure S8**. Lineage cluster membership accuracy ratio for each method using complete genomes. A ratio of 1.0 indicates perfect agreement between each method's clustering results and the TB-Profiler–assigned lineage groups. Ratios below 1.0 reflect over-clustering, where genomes from different lineage groups were incorrectly merged into the same cluster. A total of 535 genomes were included in this analysis.

**Commented [DSMR1]:** There are only for the analysis using compete genomes?

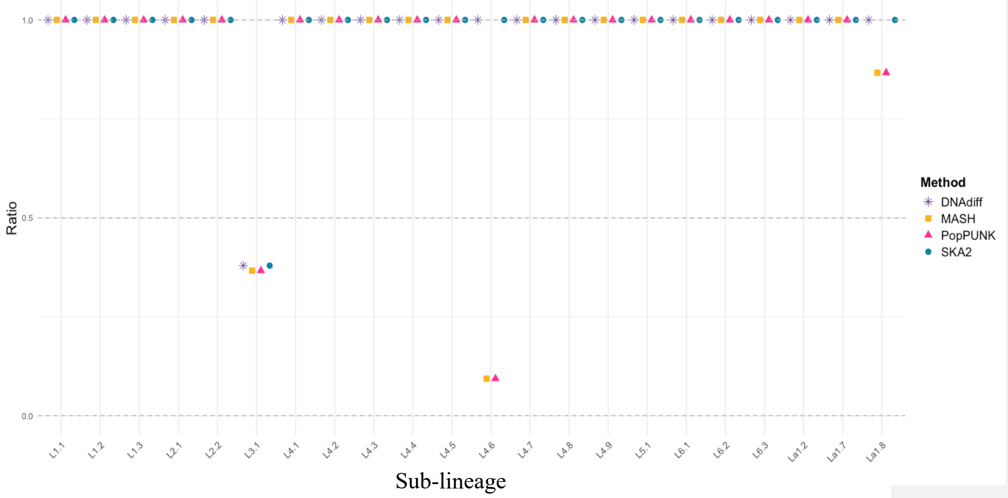


**Figure S9.** Lineage cluster membership accuracy ratio for each method using complete genomes. A ratio of 1.0 indicates perfect agreement between each method's clustering results and the TB-Profiler–assigned lineage groups. Ratios below 1.0 reflect over-clustering, where genomes from different lineage groups were incorrectly merged into the same cluster. A total of 494 genomes were included in this analysis.

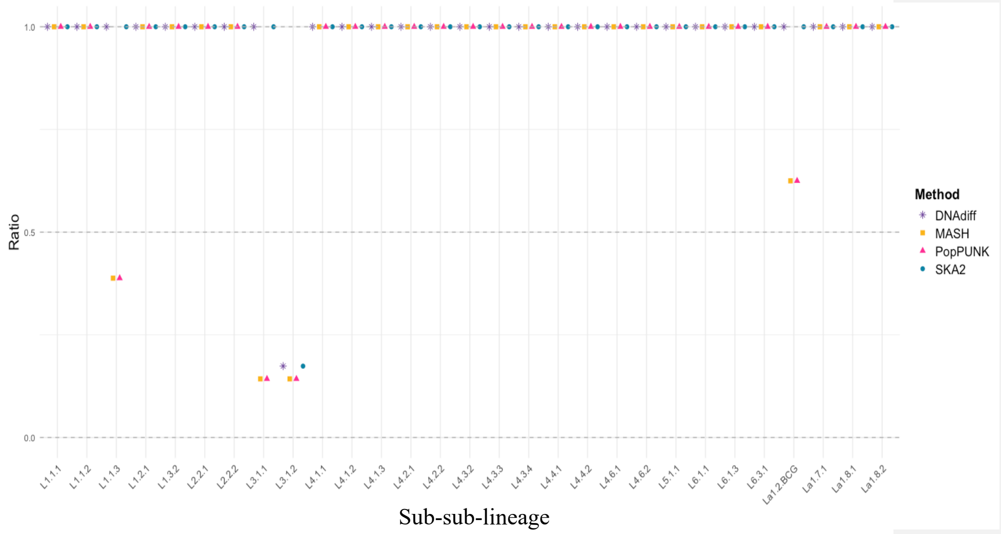


**Figure S10**. Lineage cluster membership accuracy ratio for each method using complete genomes. A ratio of 1.0 indicates perfect agreement between each method's clustering results and the TB-Profiler–assigned lineage groups. Ratios below 1.0 reflect over-clustering, where genomes from different lineage groups were incorrectly merged into the same cluster. A total of 410 genomes were included in this analysis.

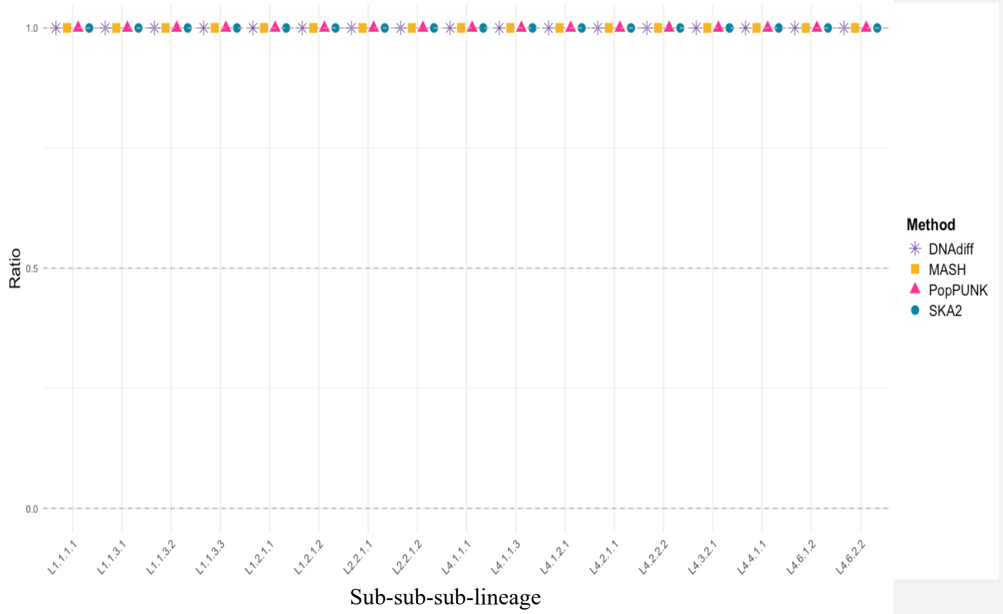


**Figure S11**. Lineage cluster membership accuracy ratio for each method using complete genomes. A ratio of 1.0 indicates perfect agreement between each method's clustering results and the TB-Profiler–assigned lineage groups. Ratios below 1.0 reflect over-clustering, where genomes from different lineage groups were incorrectly merged into the same cluster. A total of 157 genomes were included in this analysis.

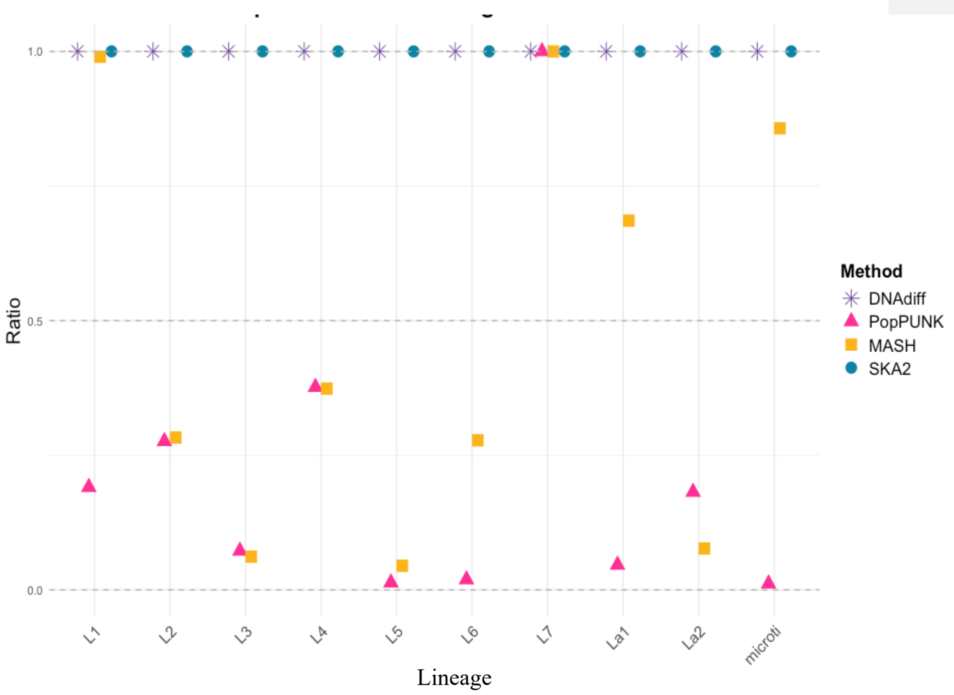


**Figure S12**. Lineage cluster membership accuracy ratio for each method using simulated reads. A ratio of 1.0 indicates perfect agreement between each method's clustering results and the TB-Profiler–assigned lineage groups. Ratios below 1.0 reflect over-clustering, where genomes from different lineage groups were incorrectly merged into the same cluster. A total of 535 genomes were included in this analysis.

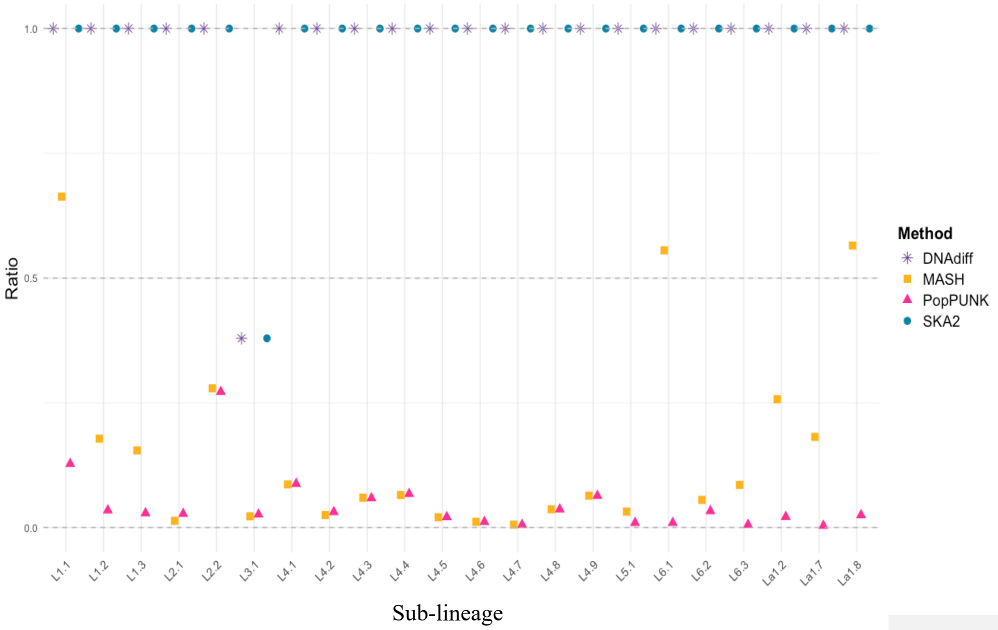


**Figure S13.** Lineage cluster membership accuracy ratio for each method using simulated reads. A ratio of 1.0 indicates perfect agreement between each method's clustering results and the TB-Profiler–assigned lineage groups. Ratios below 1.0 reflect over-clustering, where genomes from different lineage groups were incorrectly merged into the same cluster. A total of 494 genomes were included in this analysis.

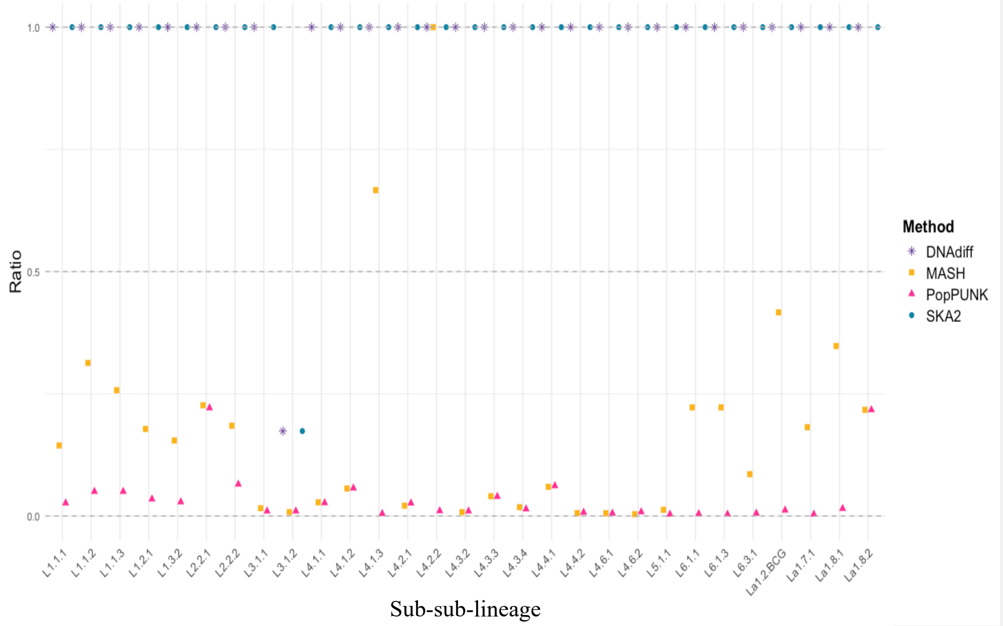


**Figure S14**. Lineage cluster membership accuracy ratio for each method using simulated reads. A ratio of 1.0 indicates perfect agreement between each method's clustering results and the TB-Profiler–assigned lineage groups. Ratios below 1.0 reflect over-clustering, where genomes from different lineage groups were incorrectly merged into the same cluster. A total of 410 genomes were included in this analysis.

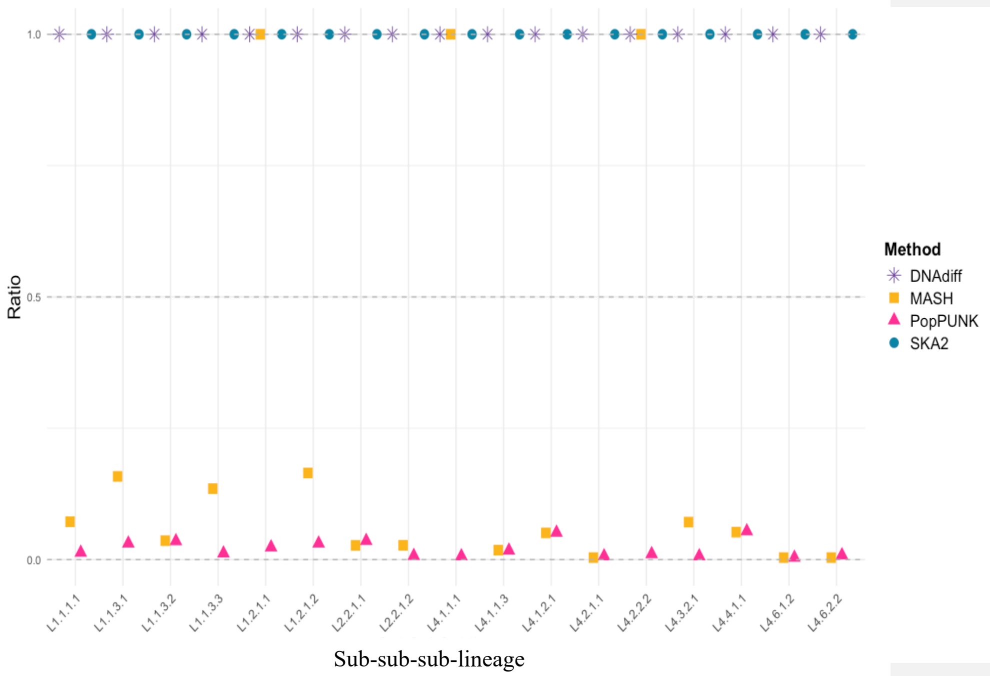


**Figure S15.** Lineage cluster membership accuracy ratio for each method using simulated reads. A ratio of 1.0 indicates perfect agreement between each method's clustering results and the TB-Profiler–assigned lineage groups. Ratios below 1.0 reflect over-clustering, where genomes from different lineage groups were incorrectly merged into the same cluster. A total of 157 genomes were included in this analysis.

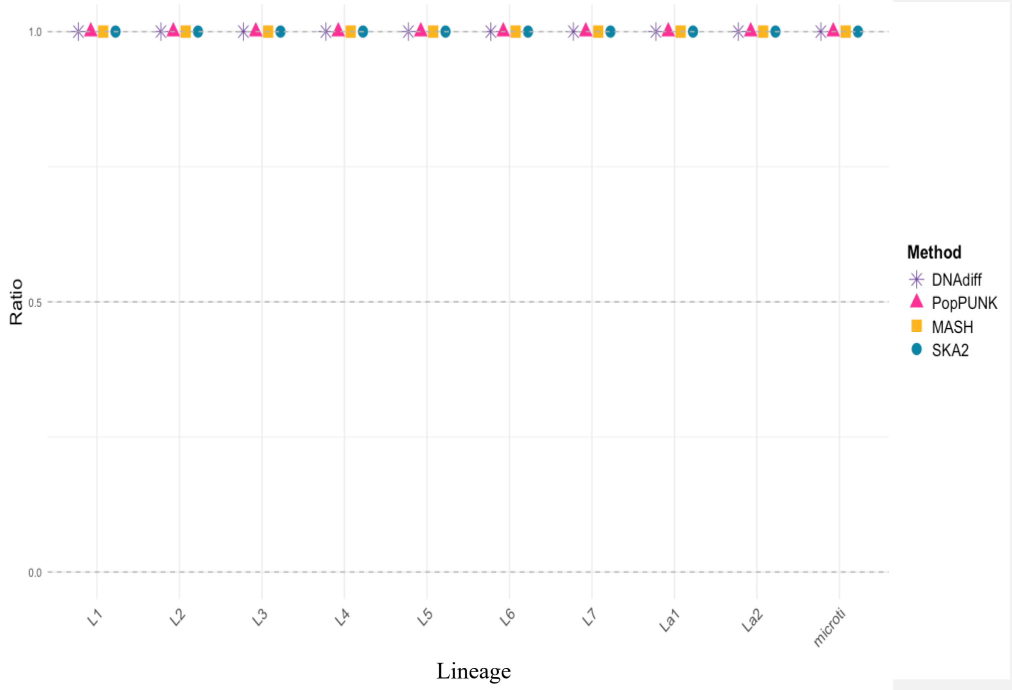


**Figure S16**. Lineage cluster membership accuracy ratio for each method using assembled simulated reads. A ratio of 1.0 indicates perfect agreement between each method's clustering results and the TB-Profiler–assigned lineage groups. Ratios below 1.0 reflect over-clustering, where genomes from different lineage groups were incorrectly merged into the same cluster. A total of 535 genomes were included in this analysis.

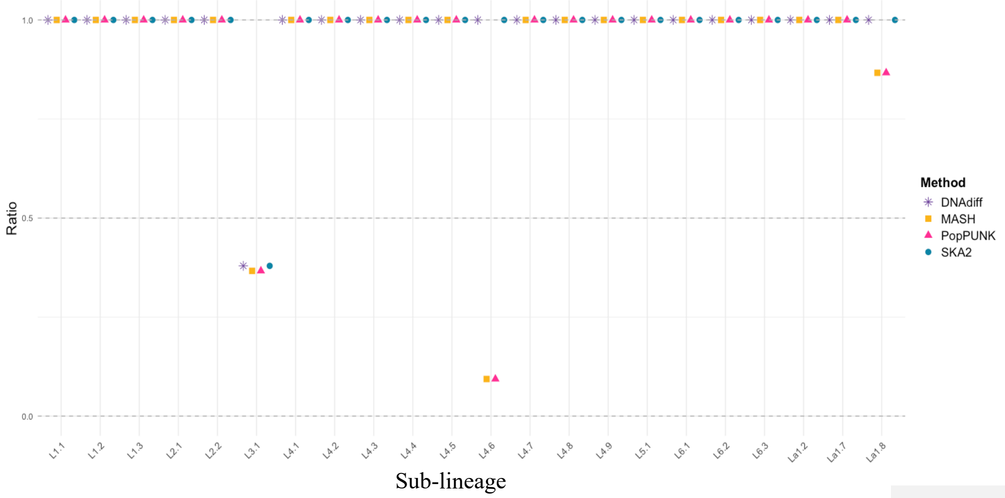


**Figure S17.** Lineage cluster membership accuracy ratio for each method using assembled simulated reads. A ratio of 1.0 indicates perfect agreement between each method's clustering results and the TB-Profiler–assigned lineage groups. Ratios below 1.0 reflect over-clustering, where genomes from different lineage groups were incorrectly merged into the same cluster. A total of 494 genomes were included in this analysis.

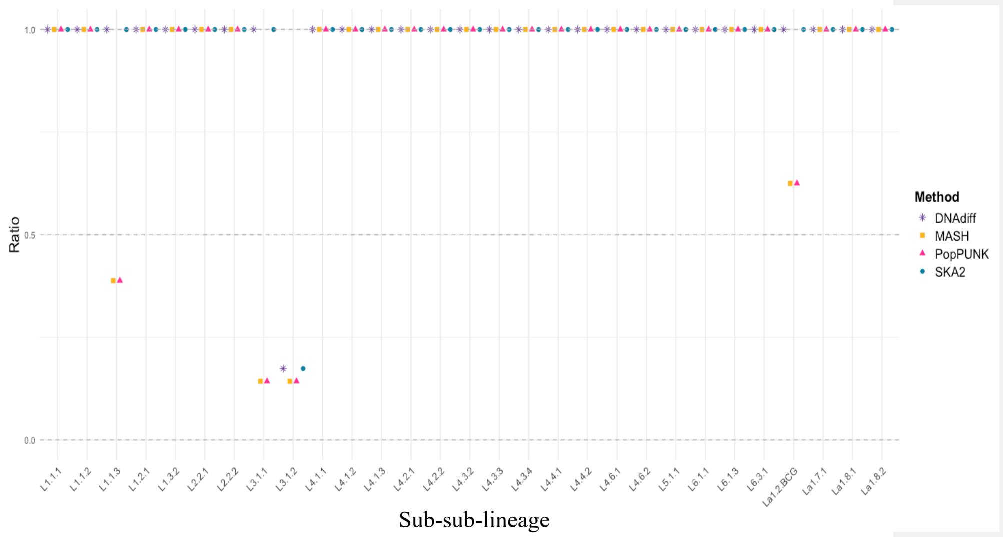


**Figure S18**. Lineage cluster membership accuracy ratio for each method using assembled simulated reads. A ratio of 1.0 indicates perfect agreement between each method's clustering results and the TB-Profiler–assigned lineage groups. Ratios below 1.0 reflect over-clustering, where genomes from different lineage groups were incorrectly merged into the same cluster. A total of 410 genomes were included in this analysis.

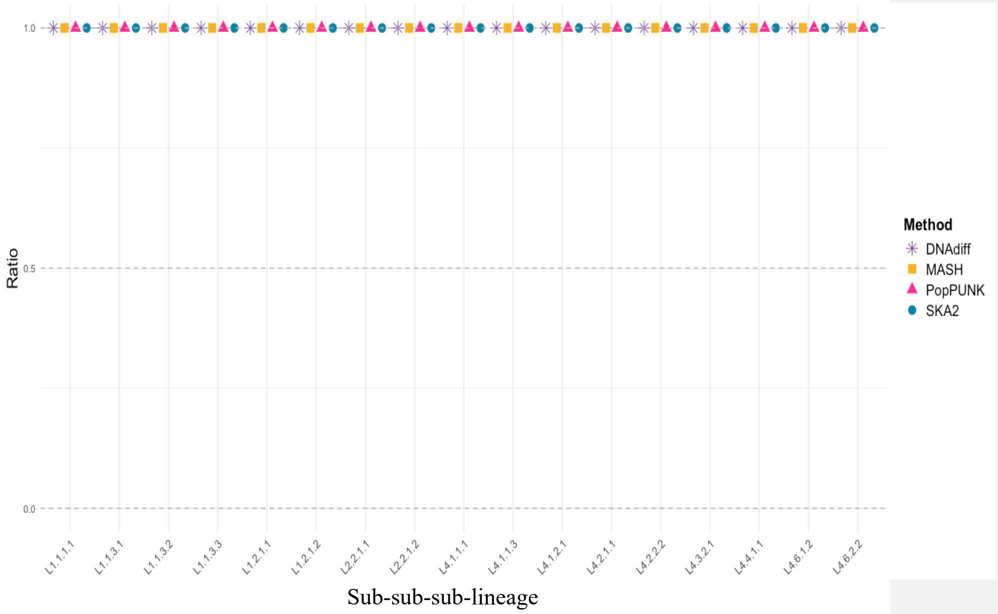


**Figure S19.** Lineage cluster membership accuracy ratio for each method using assembled simulated reads. A ratio of 1.0 indicates perfect agreement between each method's clustering results and the TB-Profiler–assigned lineage groups. Ratios below 1.0 reflect over-clustering, where genomes from different lineage groups were incorrectly merged into the same cluster. A total of 157 genomes were included in this analysis.
